# The impact of global selection on local adaptation and reproductive isolation

**DOI:** 10.1101/855320

**Authors:** Gertjan Bisschop, Derek Setter, Marina Rafajlović, Stuart J.E. Baird, Konrad Lohse

**Affiliations:** Institute of Evolutionary Biology, University of Edinburgh, UK; Department of Marine Sciences, Centre for Marine Evolutionary Biology, University of Gothenburg, Gothenburg, Sweden; Institute of Vertebrate Biology, Academy of Sciences of the Czech Republic, Brno, Czech Republic

## Abstract

Despite the homogenising effect of strong gene flow between two populations, adaptation under symmetric divergent selection pressures results in partial reproductive isolation: adaptive substitutions act as local barriers to gene flow, and if divergent selection continues unimpeded, this will result in complete reproductive isolation of the two populations, i.e. speciation. However, a key issue in framing the process of speciation as a tension between local adaptation and the homogenising force of gene flow is that the mutation process is blind to changes in the environment and therefore tends to limit adaptation. Here we investigate how globally beneficial mutations (GBMs) affect divergent local adaptation and reproductive isolation. When phenotypic divergence is finite, we show that the presence of GBMs limits local adaptation, generating a persistent genetic load at the loci which contribute to the trait under divergent selection and reducing genome-wide divergence. Furthermore, we show that while GBMs cannot prohibit the process of continuous differentiation, they induce a substantial delay in the genome-wide shutdown of gene flow.

## Introduction

Felsenstein (1981) pointed out that by any measure there are many more niches than species and demonstrated that gene flow is a strong homogenising force that will tend to prevent populations adapting to different niches. As an illustration, each broad-leaved tree provides two equal-sized niches (patches) for feeding caterpillars: under-leaf and over-leaf. These niches likely differ in predator pressure. Why then does each species not bifurcate into an under- and over-leaf phenotype? It seems they cannot, even though variation exists in nature for under- and over-leaf caterpillar lifestyles. Felsenstein’s answer is that the divergent selection caused by differing predator pressure on the pair of patches is insufficiently strong to overcome the homogenising effects of gene flow between incipient patch-populations.

Several verbal models and simulation studies have framed the process of speciation as a tension between local adaptation and the homogenising force of gene flow (Wu, 2001; Flaxman et al., 2013; Yeaman and Whitlock, 2011; Rafajlović et al., 2016). Flaxman et al. (2013) introduced BU2S (build-up-to-speciation), a model of divergent selection acting on many loci between two populations connected by gene flow. Intended as an extension of Felsenstein’s model, they concluded that their model has an emergent property: populations can adapt to the opposing environmental stresses experienced in a pair of equal-sized patches even in the face of strong gene flow and, in the process, become (partly) reproductively isolated. Multilocus extensions deriving the conditions under which a barrier with gene flow can be maintained have been described previously (e.g. Barton, 1983), and so, on superficial inspection, these results seem plausible.

The population genetics or sweep-based model simulated by Flaxman et al. (2013) assumes that local adaptation is never-ending, i.e. each new mutation confers a fixed selective advantage locally. In contrast, other simulation studies of a pair of populations connected by gene flow have taken on the quantitative genetics point of view by modeling an explicit phenotype and studying local adaptation to a new set of fixed local optima (Yeaman and Whitlock, 2011; Rafajlović et al., 2016). However, while these sweep-based and trait-based studies of divergent selection make different assumptions about the genetic basis of local adaptation (an eternal stream of local sweeps vs adaptation to a fixed set of local phenotypic optima), they share an important feature: only locally beneficial mutations (LBMs) that affect the trait(s) under divergent selection are considered. This “adaptationist” simplification ignores a central tenet of Darwinian evolution: namely, that the mutational process is blind to changes in the environment (Barton and Partridge, 2000) and therefore will tend to limit adaptation.

Ignoring deleterious mutations, a substantial fraction of new beneficial mutations must be advantageous in many environmental contexts. Returning to our toy example of caterpillars in over- and under-leaf patches, such globally beneficial mutations (GBMs) include all variants that increase caterpillar fit-ness in both leaf patches as well as any mutation that increases fitness at the adult stage. Even in the presence of a barrier to gene flow, such globally beneficial mutations will tend to selectively sweep across patches (Piálek and Barton, 1997). Previous analytic work has focused on the interaction between GBMs and LBMs in the context of adaptive introgression: how likely are GBMs to introgress from one species into another if they are linked to alleles with locally deleterious effects (Piálek and Barton, 1997; Hartfield and Otto, 2011; Uecker et al., 2015). Uecker et al. (2015) show that the probability of a single locally deleterious allele hitchhiking to fixation decays to zero over a distance 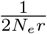, where *N*_*e*_ is the effective population size and *r* is the scaled recombination rate. For example, given parameters for modern humans (*N*_*e*_ = 10, 000 and *r* = 10^−8^), the relevant distance is ≈ 5kb, shorter than the average gene. Given that many sexual species have higher scaled recombination rates (Stapley et al., 2017), recombination should be frequent enough to prevent the majority of locally deleterious mutations from hitchhiking to fixation (Hartfield and Otto, 2011). While the sweep dynamics of individual variants that are linked to locally deleterious mutations have been characterized in some detail (Hartfield and Otto, 2011; Uecker et al., 2015), the flip-side of this interaction has received little attention: to what extent is local adaptation impeded when there is a constant supply of both GBMs and LBMs arising in a genome? Moreover, the hitchhiking-fixation of locally deleterious mutations is not necessarily the main factor slowing down adaptation. Instead, local adaptation may be limited chiefly by Hill-Robertson interference, the process whereby ongoing sweeps of mutations at partially linked loci decrease each others’ probabilities of fixation and build up negative linkage equilibrium (Hill and Robertson, 1966), between GBMs and LBMs prior to their fixation.

Here we use simulations to investigate how the presence of GBMs affects the process of divergent local adaptation and reproductive isolation. We extend the existing simulation frameworks of (Yeaman and Whitlock, 2011; Rafajlović et al., 2016; Flaxman et al., 2013) in which divergence evolves under a constant high rate of mutational influx. In these models, local adaptation involves many loci and the dynamics resulting from the selective interference of locally and globally beneficial mutations cannot be captured by the analytic results that are available for the simpler case of a single introgressing locus.

Specifically, we i) ask to what extent adaptation to locally divergent trait optima is impeded by selective sweep interference from GBMs, ii) assess how the effect of globally beneficial mutations depends on the assumptions about local adaptation (trait-based vs sweep-based models) and iii) consider how GBMs influence the evolution of reproductive isolation.

## Methods

### Trait-based model of local adaptation

We study the impact of global selection by adding globally beneficial mutations (GBMs) to a multilocus model of divergent local adaptation similar to that studied by Yeaman and Whitlock (2011) and Rafajlović et al. (2016). Simulations under this trait-based model were implemented in `SLiM3.3` (Haller and Messer, 2019). We consider two Wright-Fisher populations (discrete non-overlapping generations) of *N*_*e*_ diploid individuals that exchange, on average, *M* = 4*N*_*e*_*m* migrants per generation and experience soft selection. We considered *M* = 1 throughout, which, in the absence of GBMs, guarantees rapid local adaptation (Rafajlović et al., 2016). Recombination is modelled by assuming that cross-over events occur uniformly at random, i.e. we ignore gene conversion and physical constraints on double cross-over events. Both populations start out perfectly adapted to a shared local optimum set at 0 until the onset of divergent selection. An instantaneous change in the environment shifts their optima to *θ*_+_, and *θ_−_* = −*θ*_+_, respectively. We thus assume that adaptation is *de novo* and that the optima are stationary. We further assume that the phenotype of each individual is determined by the sum of effect sizes of all mutations affecting the trait under divergent selection. As in Rafajlović et al. (2016), the fitness of an individual in population *i* with phenotype *z* and with selection strength *σ* is given by

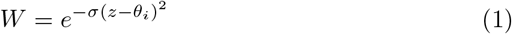

Here, *σ* denotes the strength of selection on the divergently selected trait, and is assumed to be the same in both populations. *σ* affects the width of the fitness curve with smaller values representing weak selection. Similar to Rafajlović et al. (2016) we explored two values of *σ* representing weak and strong selection on the trait under divergent selection (see Table 1). New offspring are generated each generation until the fixed population size *N*_*e*_ is reached. Each individual’s chance of being the parent to each new offspring is determined by its fitness *W*.

**Table 1:** Model parameters and values used in the trait-based model of local adaptation.

| Parameter | Explanation | Value |
| --- | --- | --- |
| $N_e$ | Size of each population | 10000 |
| $M$ | Scaled migration rate | 1 |
| $\theta$ | Optimal phenotype for the trait under divergent selection | $\theta_+ = 2, \theta_- = -2$ |
| $U$ | Genomewide mutation rate per individual | $5 \times 10^{-6}, 1.75 \times 10^{-5}, 5 \times 10^{-5}, 5 \times 10^{-4}, 1.75 \times 10^{-3}, 5 \times 10^{-3}, 5 \times 10^{-2}$ |
| $U_g/U_l$ | Ratio of the rates of GBMs to LBMs | 0, 1, 2 |
| $\alpha$ | Mean effect size of LBMs | 0.01 |
| $s_g$ | Mean of exponential distribution selection coefficients GBMs | 0.001, 0.01 |
| $\sigma$ | Overall strength of selection on LBMs | 0.0125, 0.125 |
| $p_a$ | Fraction of mutations that are beneficial | 0.01, $10^{-4}$ |
| $T$ | Total duration of the run in $2N_e$ generations | 5, 25 |
| Scenario | | $\{s_g, \sigma, p_a, T\}$ |
| weak-frequent | | $\{0.001, 0.0125, 0.01, 5\}$ |
| strong-rare | | $\{0.01, 0.125, 10^{-4}, 25\}$ |

Locally beneficial mutations (LBMs) arising at a total rate *U*_*l*_ per individual per generation, are assumed to be codominant with effect sizes drawn from a mirrored exponential distribution with mean *α*. Assuming *θ* = 2 and mean mutation effect sizes of 0.01, on average 100 mutations are required for perfect adaptation. Mutations can occur with equal probability at each position along a contiguous chromosome of map length *R*. We assume a large number of evenly spaced sites (100, 000), such that the probability of multiple mutations occurring at the same site becomes negligible (approaching an infinite sites model). If a site does get hit by a second mutation of the same type (LBM or GBM, see below), it erases the previous one (house of cards model). This differs from the model studied by Yeaman and Whitlock (2011) and Rafajlović et al. (2016) which assumes that mutations occur at a small number (50 – 2000) of evenly spaced loci with effects accumulating at loci that are hit by several mutations (continuum-of-alleles model).

### Adding globally beneficial mutations

We assume that both populations are also adapting to a shared moving optimum on a second orthogonal trait. This global optimum represents an *n*-dimensional vector co-ordinate in the space of all traits that are not under divergent selection. We assume that this trait space is so large that the optimum will never be stationary for long enough for a population to achieve perfect adaptation. To this end we further assume the population always trails this optimum at a fixed distance *θ*_+_ (the same value as for the optimum of LBMs). This makes direct comparison to the distribution of fitness effects (DFE) of LBMs possible at all times. Note however that we assume that there is no pleiotropy, i.e. each beneficial mutation contributes either to local adaptation or global adaptation, but not both. This ensures that local adaptation can be limited only by genetic hitchhiking and sweep interference of LBMs and GBMs but not pleiotropic effects of mutation on local and global fitness.

To reduce computation time we do not simulate the moving optimum explicitly, but rather draw selection coefficients for GBMs from an exponential distribution with mean *s*_*g*_. We combine the effects of LBMs and GBMs on an individual’s relative fitness in population *i* by multiplying (1) by 1 + *h*_*j*_*s*_*j*_ for all *n*_*g*_ GBMs, with *h*_*j*_ the dominance coefficient. This results in:

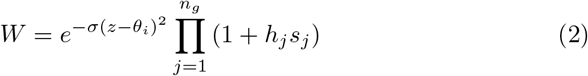

We determine *s*_*g*_ by calculating the selection coefficients for all LBMs bringing an individual with phenotype=0 (at the onset of phenotypic divergence), Δ*z* closer to *θ* (with Δ*z* drawn from an exponential distribution with mean *α*). We then match the best fitting exponential distribution to the resulting distribution of selection coefficients. Therefore, at the onset of local selection the distributions of selection coefficients of GBMs and LBMs are identical. Note however that the DFE of LBMs changes with approach to the local optima.

To measure the impact of GBMs on trait divergence, we track the mean phenotype in both populations for the trait under local selection. Phenotypic divergence is measured as the difference Δ*z* between population means, where ‘perfect’ differentiation corresponds to Δ*z* = 2*θ*_+_. We also track the time and population of origin of LBMs and use the tree sequence recording in `SLiM3.3` to record and assess the distribution of cross-population coalescence times at the end of each run (*T*) using `tskit` (Kelleher et al., 2018).

To facilitate comparisons between scenarios with and without GBMs, the rate *U*_*l*_ of LBMs is kept constant throughout. Reduced divergence is therefore not caused by a reduction in the mutational supply of LBMs. Since the relative ratio of GBMs to LBMs is unknown in nature, we explore varying *U_g_/U_l_*, but ensure that the total rate of beneficial mutations (*U*_*g*_ + *U*_*l*_ ≤ *U*) is biologically plausible given empirical estimates of *de novo* mutation rates and the distribution of fitness effects (see Choice of parameter space). For each parameter combination we ran 200 replicate simulations.

### Globally beneficial mutations in a sweep-based model

We have re-implemented the model studied by Flaxman et al. (2013) in *SLiM3* (Haller and Messer, 2019; Messer et al., 2016) by making minor changes to the trait-based model described above. For ease of comparison, we have kept model assumptions and parameters (*U*_*l*_, *U*_*g*_, *M*, *T*) the same whenever possible (Table 1). As before, we assume no pleiotropy, i.e. new mutations are either LBMs or GBMs. However, the effects of LBMs on fitness no longer depends on an explicit phenotype and a fitness function relating it to an optimum but are drawn directly from an exponential distribution with mean *s*_*l*_. Each LBM *i* confers a homozygous fitness 1 + *s*_*i*_ in population 1 and 1/(1 + *s*_*i*_) in population 2. Selection is thus purely directional. As before, co-dominance is assumed and fitness effects of all *n*_*g*_ GBMs and *n*_*l*_ LBMs (at any point in time) across multiple loci are multiplicative. Selection coefficients for GBMs and LBMs are drawn from the same exponential distribution. The impact of GBMs on fitness is modelled in the same way as before resulting in:

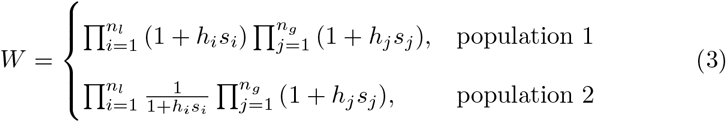

### Choice of parameter space

We have sought to cover a biologically plausible range of parameters including humans and *Drosophila*. We focus on two scenarios which are loosely motivated by two contrasting estimates of the fraction of beneficial mutations and the DFE in *Drosophila melanogaster*: Schneider et al. (2011), using inferences based on the site frequency spectra of synonymous and nonsynonymous nucleotide sites, estimated that a fraction *p*_*a*_ = 0.01 of mutations is selectively advantageous with a mean effect size of *N*_*e*_*s* = 5. In contrast, recent estimates based on the correlation between within-population synonymous nucleotide site diversity of a gene and its divergence from related species at nonsynonymous sites suggest that beneficial mutations may be much rarer (*p*_*a*_ = 10^−4^) (Campos et al., 2017) but of larger effect (mean *N*_*e*_*s* = 250). Given that both estimates are likely biased by their contrasting assumptions about the effect of selection on linked neutral sites and the fact that targets of selections are unlikely to be uniformly distributed in real genomes, we have implemented both a “weak-frequent” (weak selection, frequent mutations) and a “strong-rare” scenario and explored a range of *U* values for each case (see Table 1 for parameter values).

All simulations were run for two different lengths of sequence *R* = 0.5 and 0.05 Morgan. Assuming a typical *Drosophila* chromosome (*R* = 0.5M) of length 50Mb, a rate of *de novo* mutation *μ* = 3.5 · 10^−9^ per base and generation Keightley et al. (2009) and *p*_*a*_ = 0.01 (i.e. the weak-frequent scenario), we expect a total rate of beneficial mutations *U* = 0.00175 per generation.

### Data availability

All scripts are available at https://github.com/GertjanBisschop/GBMs.

## Results and Discussion

The results section is structured as follows: we first describe the effect of GBMs on adaptation to a fixed set of local trait optima. We then ask how GBMs affect genome-wide divergence measured in terms of the distribution of between population coalescence times and consider their influence on clustering of LBMs. Finally, we investigate GBMs in the context of the more extreme model of continued divergence considered by Flaxman et al. (2013) and discuss their effect on the evolution of reproductive isolation during speciation. We focus primarily on the results for the larger map length (0.5 Morgan) which is biologically more realistic (analogous to one chromosome).

### GBMs reduce local adaptation

Comparing the mean divergence along the local trait axis between simulations with and without GBMs shows clearly that GBMs reduce mean phenotypic divergence, and thus local adaptation (Δ*z* in figure 1). Unsurprisingly, the effect of GBMs on local adaptation depends on the relative mutation rates of GBMs to LBMs (*U*_*g*_/*U*_*l*_, yellow vs orange in figure 1). We do not find any qualitative differences between the “strong-rare” and the “weak-frequent” scenario.

**Figure 1:**
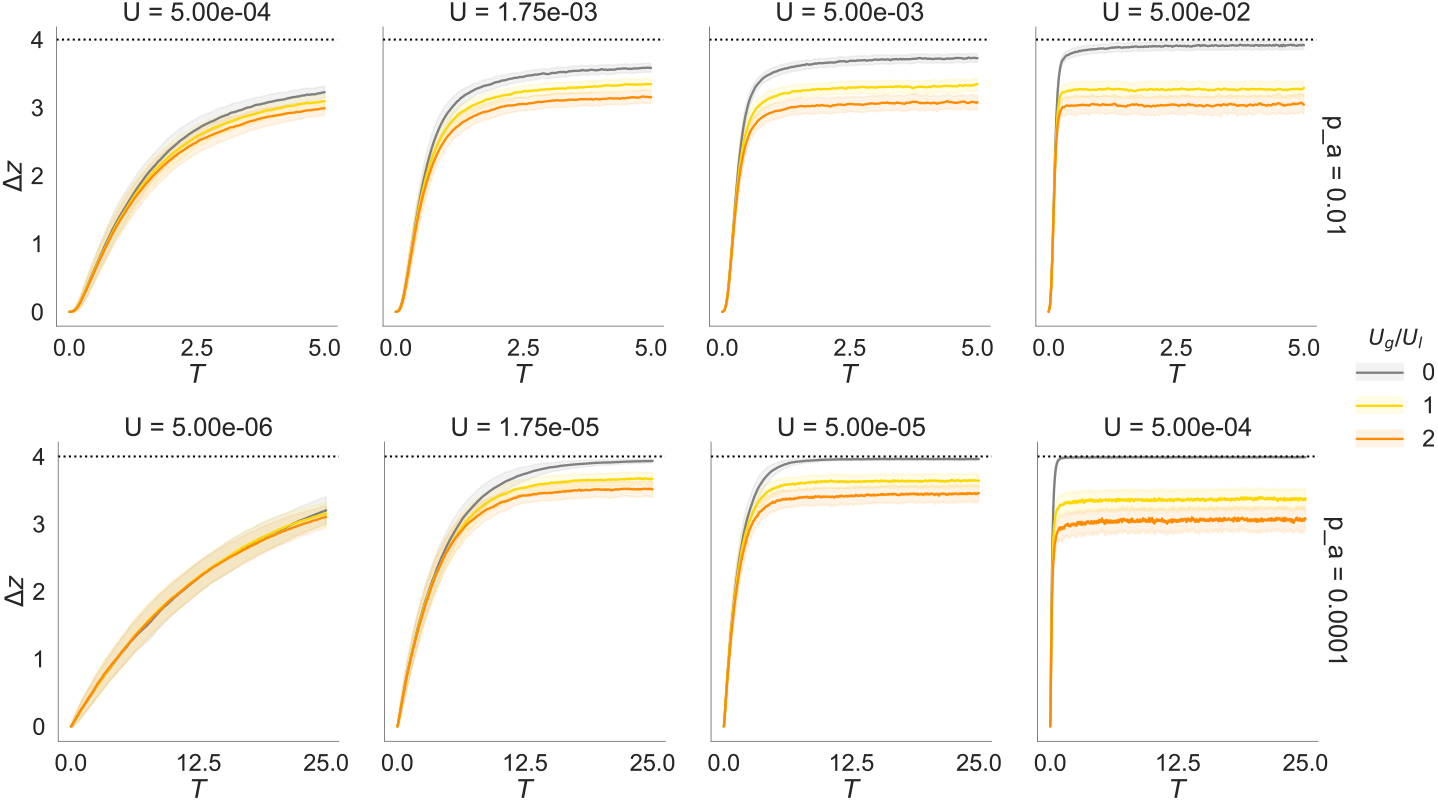
Mean trait divergence across 200 replicates (the maximum divergence is Δ*z* = 4) without GBMs (gray) and with GBMs (*U*_*g*_/*U*_*l*_ ∈ {1, 2}, coloured lines). Top row: the “weak-frequent” (*p*_*a*_ = 0.01) scenario. Bottom row: the “strong-rare” (*p*_*a*_ = 1 × 10^−4^) scenario. The time (in 2*N*_*e*_ generations) it takes for two populations to diverge in average phenotype and the effect of GBMs depends on the total rate of beneficial mutations *U*. Since the map length *R* = 50*cM* is fixed, the mutational input and the number of GBMs sweeping at any given time increases from left to right, i.e. both adaptation and the impact of GBMs increase with *U*. The envelopes show 2 standard deviations across replicates.

Rather than slowing down local adaptation, GBMs halt the process at a certain point, thus decreasing the (maximum) possible mean trait divergence between populations. In other words, GBMs induce a constant genetic load (Lenormand, 2002) by reducing the proportion of locally beneficial LBMs segregating in the population and increasing the proportion of locally maladaptive LBMs (figure 2). Although the mean trait divergence asymptotes to an equilibrium value below 2*θ*_+_, we want to emphasize that this equilibrium is highly dynamic, i.e. subject to high variance both within and among individual runs (figures 1 and 3). For large map lengths (0.5 M), GBMs do not substantially impede local adaptation, but lead to more erratic evolutionary trajectories. Assuming a shorter map length (0.05 M), i.e. GBMs arise in much closer association to LBMs, leads to a more pronounced reduction of local adaptation (figure S1).

**Figure 2:**
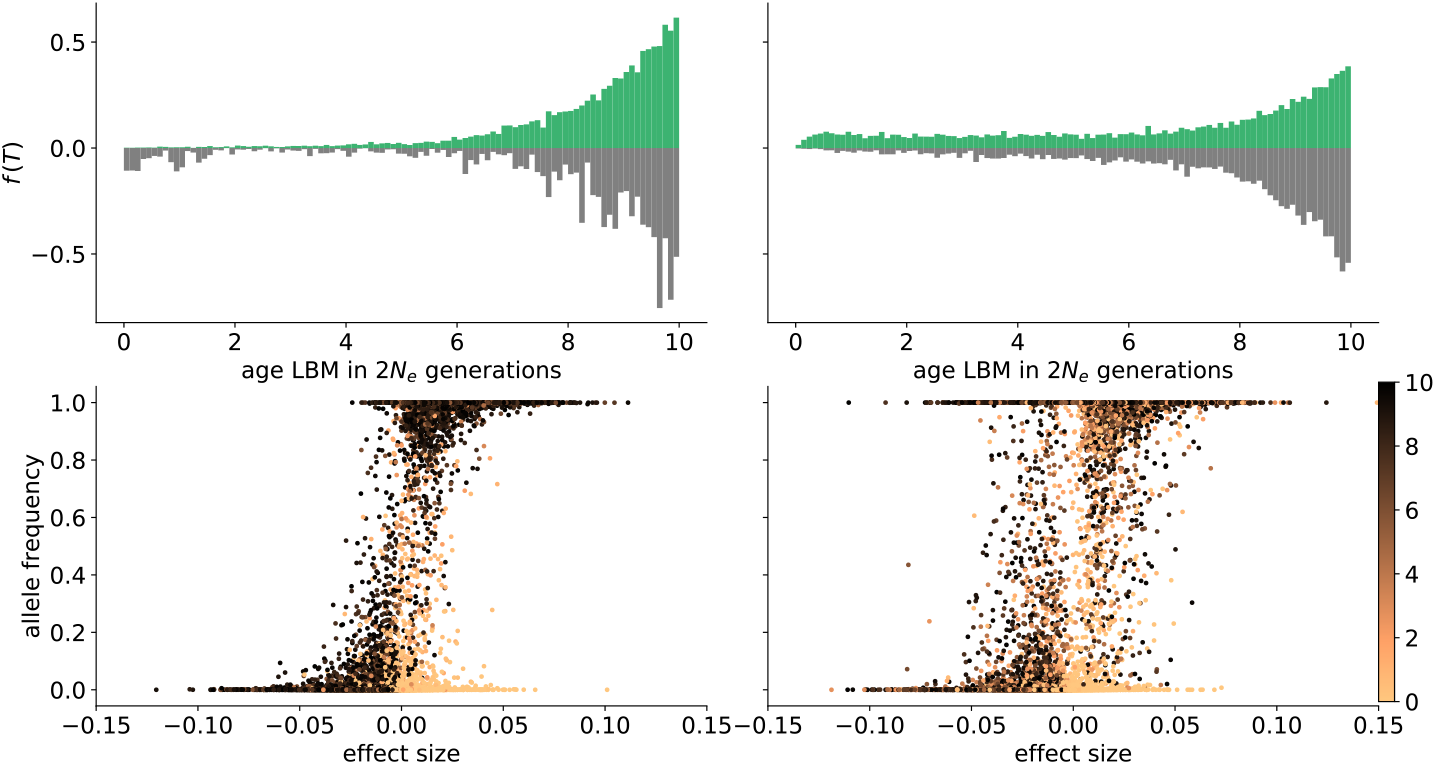
The impact of GBMs on the age distribution of LBMs. Results without GBMs (left) and with GBMs (*U*_*g*_/*U*_*l*_ = 2, right) for “the strong-rare” scenario, for the population adapting to *θ*_+_, at the dynamic equilibrium phase (10 × 2*N*_*e*_ generations). Top row: the age distribution of LBMs (across the genome and across 200 replicate runs) weighted by their frequency and effect size. Age is measured in 2*N*_*e*_ generation with 0 representing the time of sampling. The histograms on the top (green) and bottom (grey) correspond to LBMs with positive and negative local effects respectively. Bottom row: LBM effect size plotted against the allele frequency. Young and old LBMs grade from orange to black respectively (see color bar).

**Figure 3:**
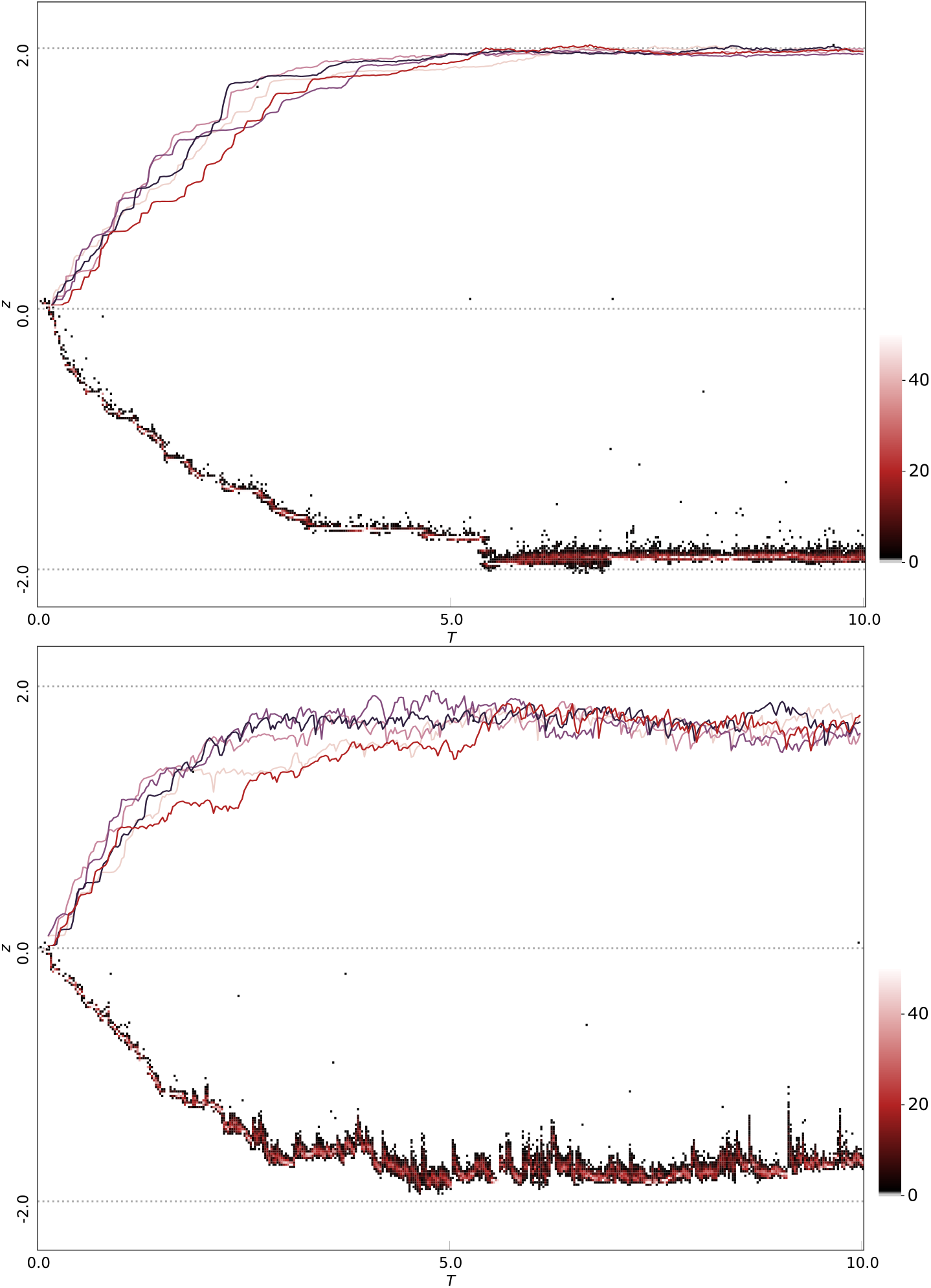
The mean phenotype through time for five simulation replicates (top half of each graph) and the evolution of individual trait values *z* for a single run (bottom half) for 10 × 2*N*_*e*_ generations for the “strong-rare” scenario. The local optimum is represented by the dotted line (*θ*_−_ = −2 and *θ*_+_ = 2). Top panel: without GBMs. Bottom panel: with GBMs (*U_g_/U_l_* = 2 and *U* = 5 × 10^−5^. The bottom half of each panel shows the distribution of phenotypes in 100 individuals (sampled every 500 generations). The shade of each dot represents the number of individuals per bin (see colorbar). 15Data corresponds to iteration(s) from figure 1 panel (2,3).

To understand why the observed dynamics differ from the impact of single introgression events of beneficial mutations (here GBMs) associated with a deleterious load (here foreign LBMs), it is useful to consider the continuous process of sweeping GBMs during the adaptive trajectory of a population: at the onset of divergent selection, each population is displaced far from the new local optimum. As a consequence, the initial local adaptation is due to a few mutations of large effect that rapidly sweep to fixation (Orr, 1998). During this phase, which involves a small number of rapid sweeps of large effect LBMs, segregating GBMs are unlikely to occur in close proximity to these LBMs.

The subsequent approach to the optimum proceeds through mutations of progressively smaller effect sizes (figure 3) with longer sojourn times. Given the increasing number of segregating LBMs, the probability that GBMs arise in close proximity increases, as does the chance of deleterious LBMs rising to appreciable frequencies before they are dissociated from sweeping GBMs by recombination (figure 3). As the fitness landscape becomes less steep upon approaching the optimum, the time it takes and the frequency at which locally deleterious LBMs recombine away from sweeping GBMs is increased (Uecker et al., 2015). Once a LBM is at appreciable frequency, its trajectory will be affected by GBMs that arise sufficiently close (and in the same population).

To summarise, the effects of sweeps on the adaptive trajectories of populations overlap (in time) for many generations. Thus analytical results for sweeping GBMs occurring in isolation from each other (single introgression events) cannot capture the potential sweep interference between LBMs and GBMs. For example, although the frequency trajectory of an established LBM can only be affected by GBMs in a small fraction of genome around it (Hartfield and Otto, 2011; Uecker et al., 2015), introgression happens continuously. Even if deleterious LBMs recombine away and are purged, their sojourn time in the ‘wrong’ population will be increased relative to a scenario without GBMs (figure 2). Sweep interference results from the time overlap of these sojourns, and its potential to limit adaptation is increased with GBMs. Additionally, while the probability of any individual GBMs dragging locally deleterious LBMs to fixation is low, such events do accumulate (figure 2 right, bottom panel) and are compensated by new LBMs. This turn-over is visible in the age distribution of LBMs which, in the presence of GBMs, shows a skew towards the recent past (figure 2 top). In contrast, without GBMs, LBMs contributing to local adaptation tend to be old, i.e. have fixed during the initial phase of local adaptation (figure 2, left).

### GBMs reduce genome wide divergence and increase clustering

The evolution of reproductive isolation can be characterized as a gradual reduction in effective migration rate (*m*_*e*_) (Barton and Bengtsson, 1986). A measurable (in practice) metric for long-term *m*_*e*_ is the genome-wide distribution of pairwise coalescence times between species, *f*(*T*_2_). As long as gene flow is ongoing, recent coalescence is likely. However, once reproductive isolation is complete, *f*(*T*_2_) is fixed and can only shift pastwards. Inspection of *f*(*T*_2_) genome-wide (averaged across simulation replicates) shows that in the absence of GBMs, selection against maladapted migrants reduces *m*_*e*_ (Aeschbacher et al., 2017) and hence the chance of recent coalescence compared to the neutral expectation (figures 4 and S6). In contrast, the continued global sweeps induced by introgressing GBMs have the opposite effect of increasing *m*_*e*_ and the fraction of genome with recent ancestry.

**Figure 4:**
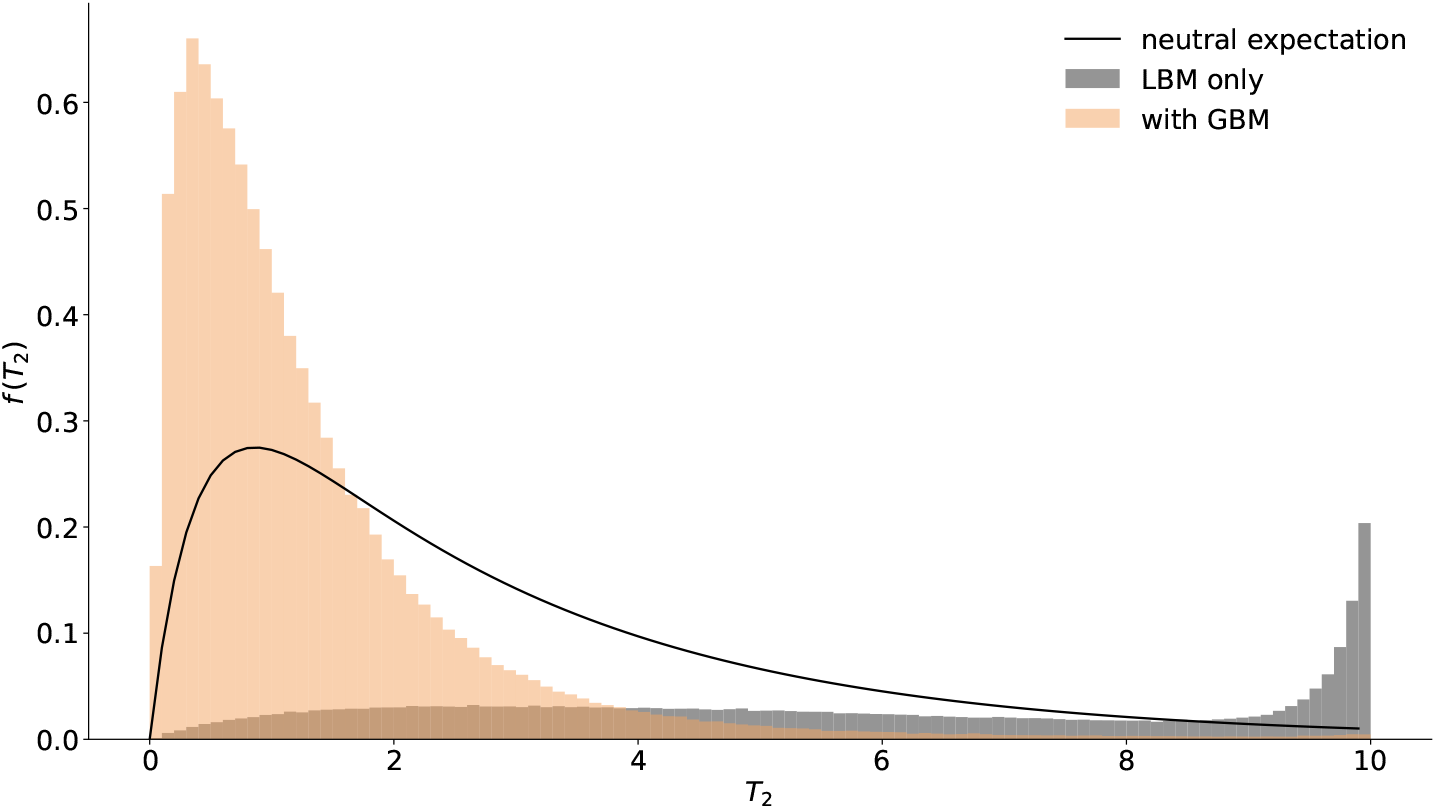
The genome-wide distribution of pairwise coalescence times *f*(*T*_2_) for the “strong-rare” scenario. The neutral expectation (assuming *M* = 1) (Lohse et al., 2016, eq. 10) is shown as a black solid line, distributions in the presence (*U*_*g*_/*U*_*l*_ = 2) and absence of GBMs are shown in tan and gray respectively. 69% (without GBMs) and 99% (with GBMs) of pairwise coalescence times are smaller then 10 × 2*N*_*e*_ generations. Data corresponds to figure 1 panel (2,3).

The question of whether local adaptation in the face of gene flow leads to clustered genetic architectures for local adaptation has received much attention (Yeaman and Whitlock, 2011; Rafajlović et al., 2016). Given that Yeaman and Whitlock (2011) described weak selection promoting clustering, we compared the distribution of pairwise distances between consecutive LBMs in the presence and absence of GBMs for the “weak-frequent” scenario. Similar to Yeaman and Whitlock (2011) we conditioned on LBMs that contribute to phenotypic divergence. Summarizing this distribution across simulation replicates and weighing LBMs by their effect size shows that the presence of GBMs increases clustering (figure S7). However, we stress that even over the long timespan we consider, both the amount of clustering that emerges overall and the increase in clustering due to GBMs are small. While our results confirm previous findings that clustering emerges even in the absence of GBMs, a direct comparison with Yeaman and Whitlock (2011) and Rafajlović et al. (2016) is challenging given the differences in models: while we approximate an infinite sites model, Yeaman and Whitlock (2011) and Rafajlović et al. (2016) assume a limited number of loci at which mutational effects build up. Thus their model has clustering inbuilt and it is perhaps unsurprising that, given enough time, large effect loci arise (Yeaman and Whitlock, 2011, figure 4).

Booker et al. (2019) have recently shown that GBMs also present a challenge for attempts to infer the targets of divergent selection in genome scans and may be impossible to distinguish from LBMs using *F*_*st*_ (Charlesworth, 1998; Cruickshank and Hahn, 2014). However, given the different effects LBMs and GBMs have on the distribution of between population coalescence times, it should be possible to distinguish them in real data using richer summaries of sequence variation than *F*_*st*_. Indeed, powerful methods for detecting individual introgression sweeps already exist (Setter et al., 2019). An interesting direction for future work will be to exploit the information contained in the distribution in coalescence times around a large number of putative targets of local and global selection in genome-wide data. This can only increase the power for such inference.

### GBMs delay the build up of strong reproductive isolation

We have so far only considered the effect of GBMs on adaptation to a fixed set of local optima. The scenario of recurrent sweeps of LBMs considered by Flaxman et al. (2013) is both simpler, more extreme (and arguably less realistic). Flaxman et al. (2013) do not consider phenotypes explicitly but instead assume that local selection is never-ending, which is equivalent to assuming eternally diverging local optima. While in this case strong reproductive isolation (“genome congealing”) is inevitable, GBMs still affect the speed at which reproductive isolation evolves. Although GBMs do not result in any increase in short term *m*_*e*_ measured over a single generation as the relative average fitness of migrants (figure S8), their long-term cumulative effect is to delay the completion of strong reproductive isolation. This delay in the genome-wide shutdown of gene flow is reflected in the widening of the distribution of between-population coalescence times (figure S9).

### Conclusions and future directions

We set out to address an ‘adaptationist’ imbalance in existing models of evolution under divergent selection. We believe there is no reason to assume that following the onset of local adaptation, all beneficial mutations will be subject to divergent selection. We termed the remaining fraction globally beneficial mutations (GBMs) and studied their impact on local adaptation and the emergence of reproductive isolation under two existing models.

We show that GBMs lead to more erratic evolutionary trajectories during local adaptation in the face of gene flow and may slow down the evolution of reproductive isolation regardless of whether we consider local adaptation to a fixed set of optima or as a runaway process. Our implementation approximates an infinite-sites-model with uniformly distributed mutational targets for both LBMs and GBMs. Given the importance of linkage, any deviation from our assumption of a polygenic trait under divergent selection, or from the assumption that mutational targets for LBMs and GBMs do not arise in strictly seperate parts of the genome, will reduce the impact of GBMs on local adaptation. We only considered *U*_*g*_/*U*_*l*_ of up to 2. While the ratio of GBMs to LBMs is an unknown quantity in nature (again depending on the distribution and the number of potential sites of LBMs and GBMs relative to each other), it seems plausible that there are many more mutations that affect global fitness than local adaptation.

Our intent was to explore a more general mutational model for adaptation under divergent selection. However, for the sake of comparability, we have followed many of the same simplifying assumptions as existing work, i.e. complete symmetry between populations both in terms of their demography (*N*_*e*_, *M*) and selection (an instantaneous switch to symmetric trait optima). We have also introduced our own symmetric simplification: the same DFE for GBMs and LBMs. Given that such many-fold symmetry is biologically unrealistic, an important task for future work is to test the robustness of these results to violations of symmetry. While we can venture to make educated guesses about relaxing symmetry assumptions for individual parameters (e.g. GBMs should reduce local adaptation in both populations even when only one is given a new optimum), the effects of GBMs under a completely general model are difficult to intuit. Further research with these kinds of models would benefit from studying the impact of hard selection, given that these models rely on continuous diverging selection for very long time periods. Secondly, it would be helpful to understand to what extent GBMs can facilitate the local fixation of recombination modifiers such as inversions (Navarro and Barton, 2003) or chromosomal fusions.

The theme of this special issue is the evolution of strong reproductive isolation during speciation. It might be suggested that our results are of little relevance since they appear to address primarily the initial evolution of reproductive barriers through adaptation to stable but divergent trait optima in two populations connected through migration. We show that in this case, migration shuts down only in regions of the genome which contribute to local adaptation causing partial reproductive isolation. However, we would argue that the the equilibrium shown in figure 1 corresponds to a stable endpoint, despite the fact that barriers between locally adapted taxa remain permeable and globally beneficial genes can introgress with negligible delay (Piálek and Barton, 1997). Biologists studying gene flow in nature discarded the biological species concept (BSC) more than a quarter of a century ago (Mallet, 1995) because its simplistic requirement of complete reproductive isolation, while attractive for the purposes of categorisation, contradicts the widespread evidence of hybridisation. There is now strong evidence from genomic data that species barriers have been permeable across entire radiations (e.g. *Heliconius* butterflies (Edelman et al., 2019)). Likewise, the discovery that humans have received adaptive introgression across permeable species barriers from archaic hominins (Racimo et al., 2015) has not led to a re-categorisation of hominins as a single BSC species due to their incomplete reproductive isolation.

Taxa that maintain distinctive genomes in the face of gene flow are separated by permeable reproductive barriers. If maintenance is exogenous, for example depending on niche existence, environmental change will make them transient (Seehausen et al., 2008). If endogeneous barriers develop, long-term persistence becomes more likely. In this context, it would be interesting to test how incorporating epistasis between LBMs (for example by modifying the exponent in the fitness function (Fraïsse et al., 2016)) in our model would affect the evolution of reproductive isolation. Although if we assume adaptation to be highly polygenic, epsistasis is likely to only have modest effects (Barton, 2017).

While we tend to focus on speciation as the evolution of complete isolation and so-called congealing of the genome, partial reproductive isolation may be an alternative and evolutionary stable endpoint, even in the presence of other mechanisms that reduce genetic exchange between locally adapted populations such as epistasis and reinforcement (Servedio and Hermisson, 2019).

GaBaM!

## Acknowledgements

We thank Nick Barton and Jarrod Hadfield for discussions, Tom Booker for constructive criticism of the manuscript. This work was supported by an ERC starting grant (ModelGenomLand). KL was supported by a NERC fellowship (NE/L011522/1). SJEB was supported by GACR grant 17-25320S. MR was supported by the Hasselblad Grant for Female Scientists, and by grants from Swedish Research Councils (Formas and VR) to the Centre for Marine Evolutionary Biology at the University of Gothenburg (www.cemeb.science.gu.se). This work has made use of the resources provided by the Edinburgh Compute and Data Facility (ECDF) (http://www.ecdf.ed.ac.uk/).

## Supplementary information

**Additional figures trait-based model**

**Additional figures sweep-based model**

**Figure S1:**
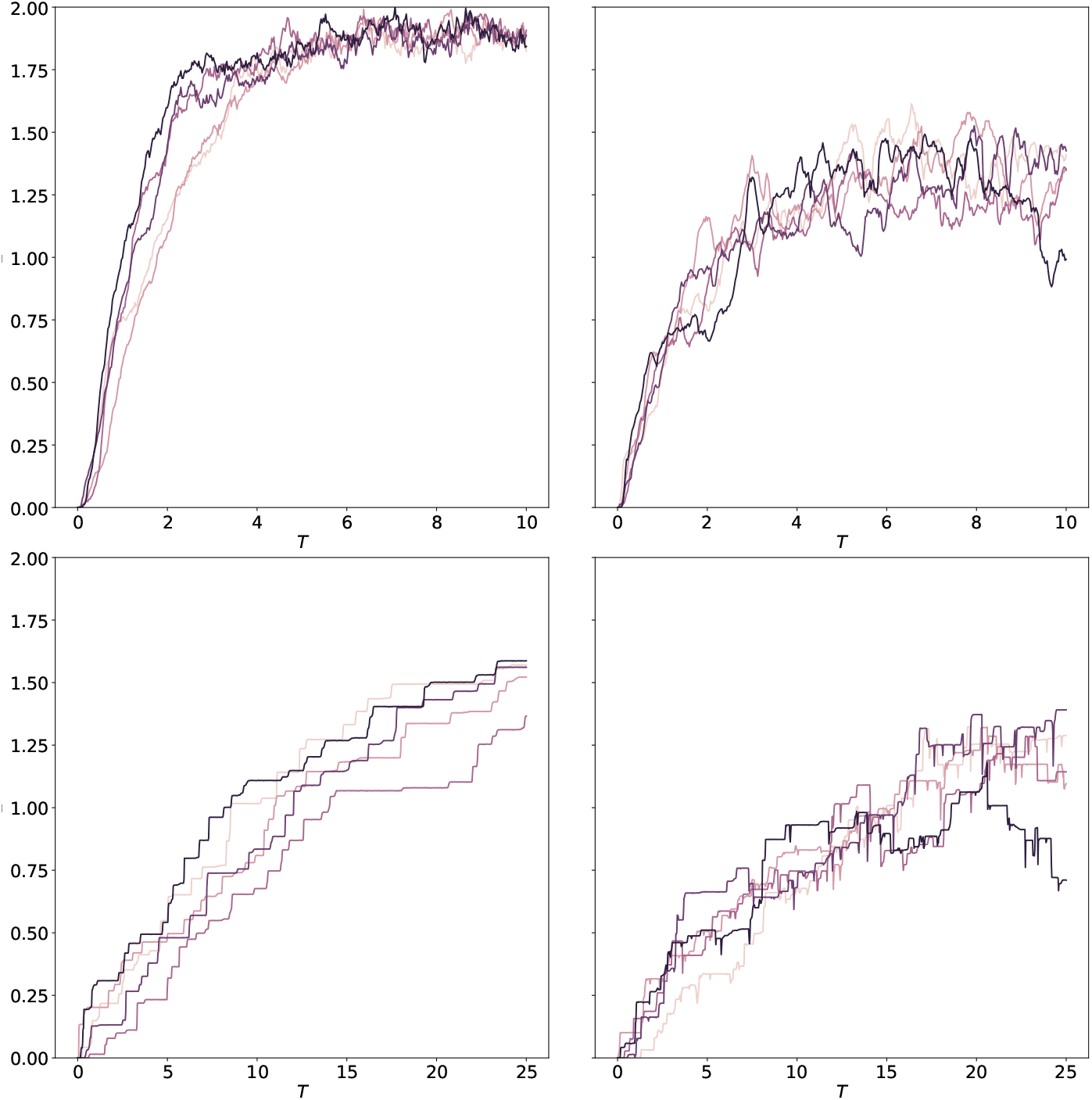
Left: LBMs only, Right: with GBM. Top row: “weak-frequent”. Bottom row: “strong-rare”. Mean phenotype for the population adapting to *θ*_+_, six replicates are shown for each scenario.

**Figure S2:**
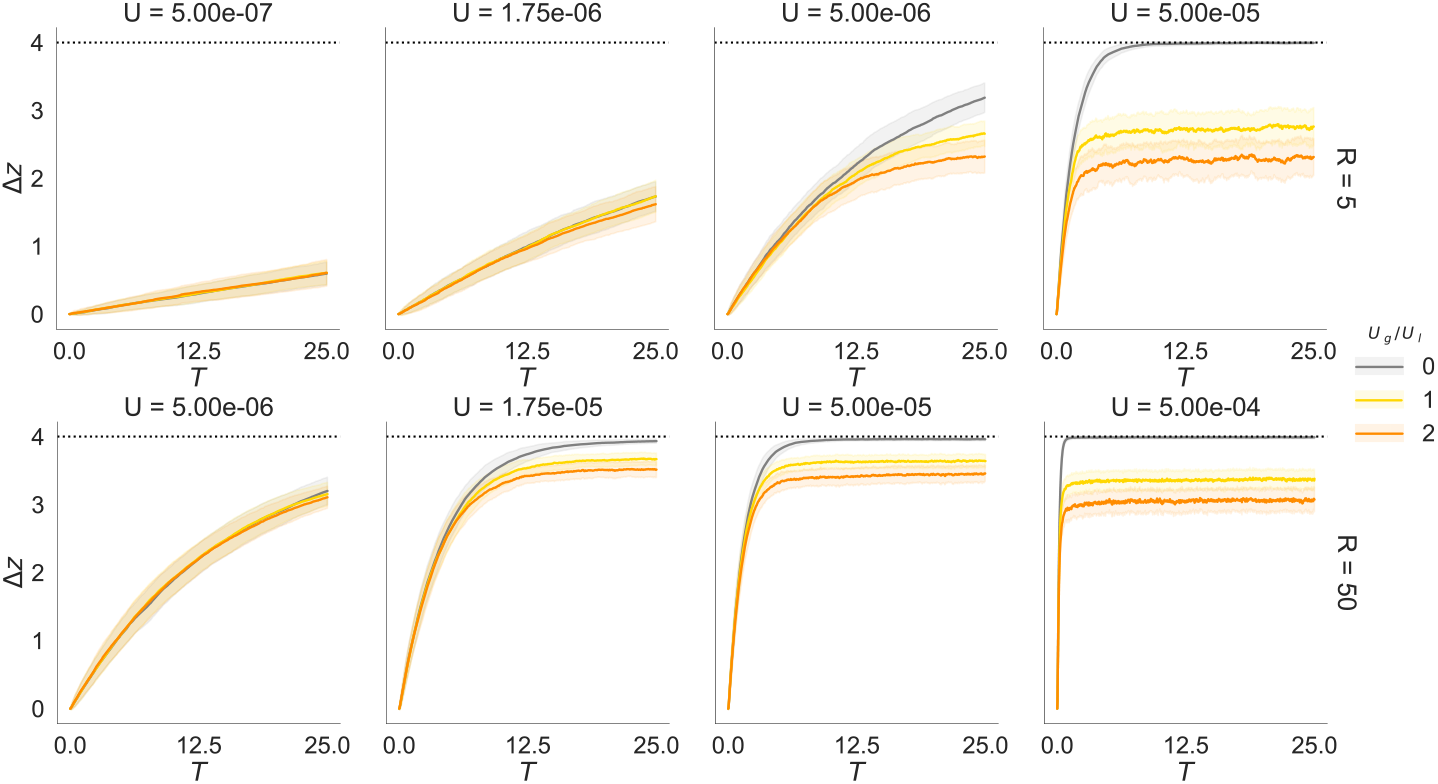
Mean trait divergence (Δ*z*) across 200 replicates (maximum Δ*z* = 4) for the “strong-rare” case without GBMs (gray) and with GBMs (*U*_*g*_/*U*_*l*_ ∈ {1, 2}, coloured lines). The envelopes show 2 standard deviations across replicates (Top *R* = 5*cM*, bottom *R* = 50*cM*). We have adapted the mutation rate assuming that the mutational input for an organism with the same average mutation and recombination rate is smaller for a smaller chromosome fragment (*R* = 50*cM* vs *R* = 5*cM*).

**Figure S3:**
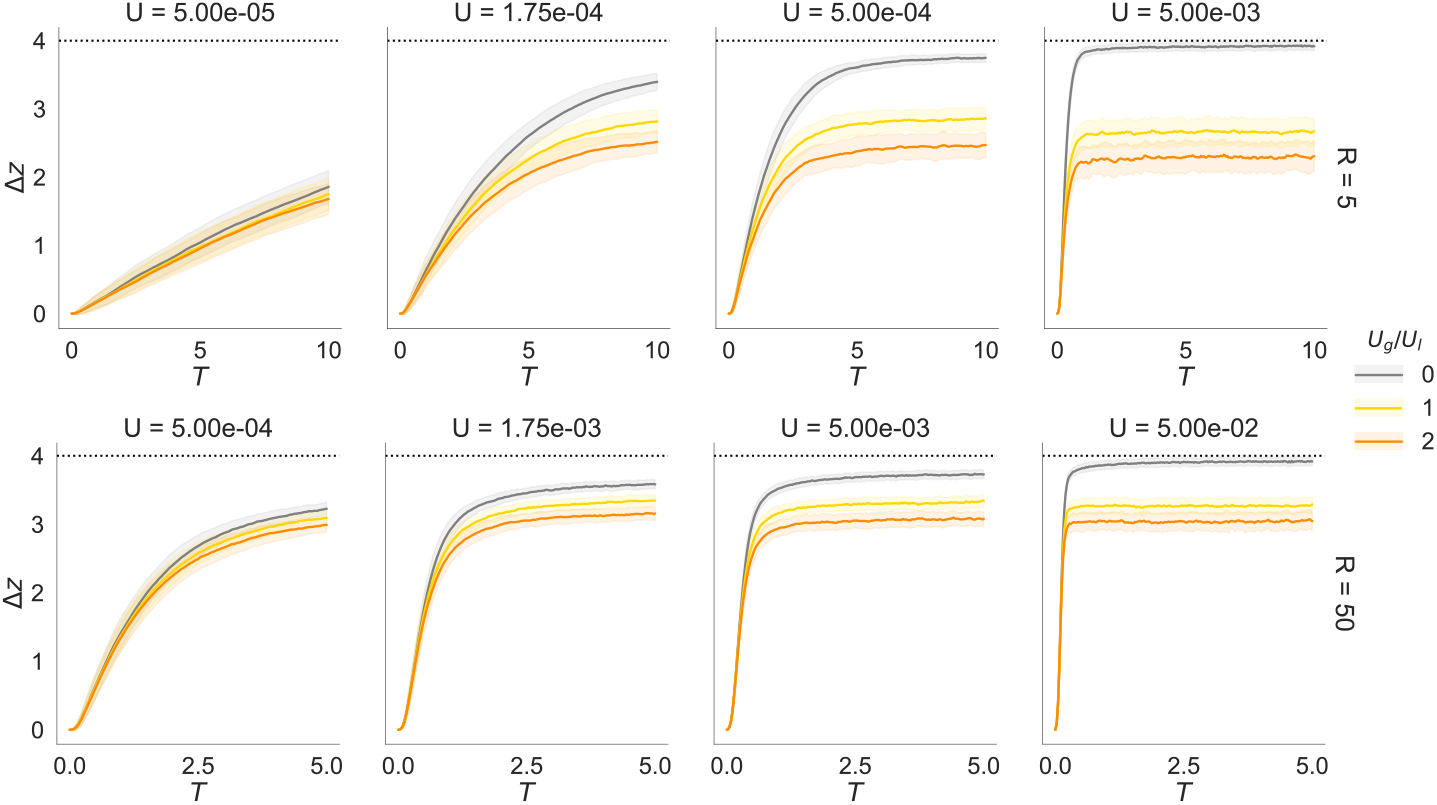
Mean trait divergence (Δ*z*) across 200 replicates (maximum Δ*z* = 4) for the “weak-frequent” case without GBMs (gray) and with GBMs (*U*_*g*_/*U*_*l*_∈ {1, 2}, coloured lines). The envelopes show 2 standard deviations across replicates (Top *R* = 5*cM*, bottom *R* = 50*cM*). We have adapted the mutation rate assuming that the mutational input for an organism with the same average mutation and recombination rate is smaller for a smaller chromosome fragment (*R* = 50*cM* vs *R* = 5*cM*).

**Figure S4:**
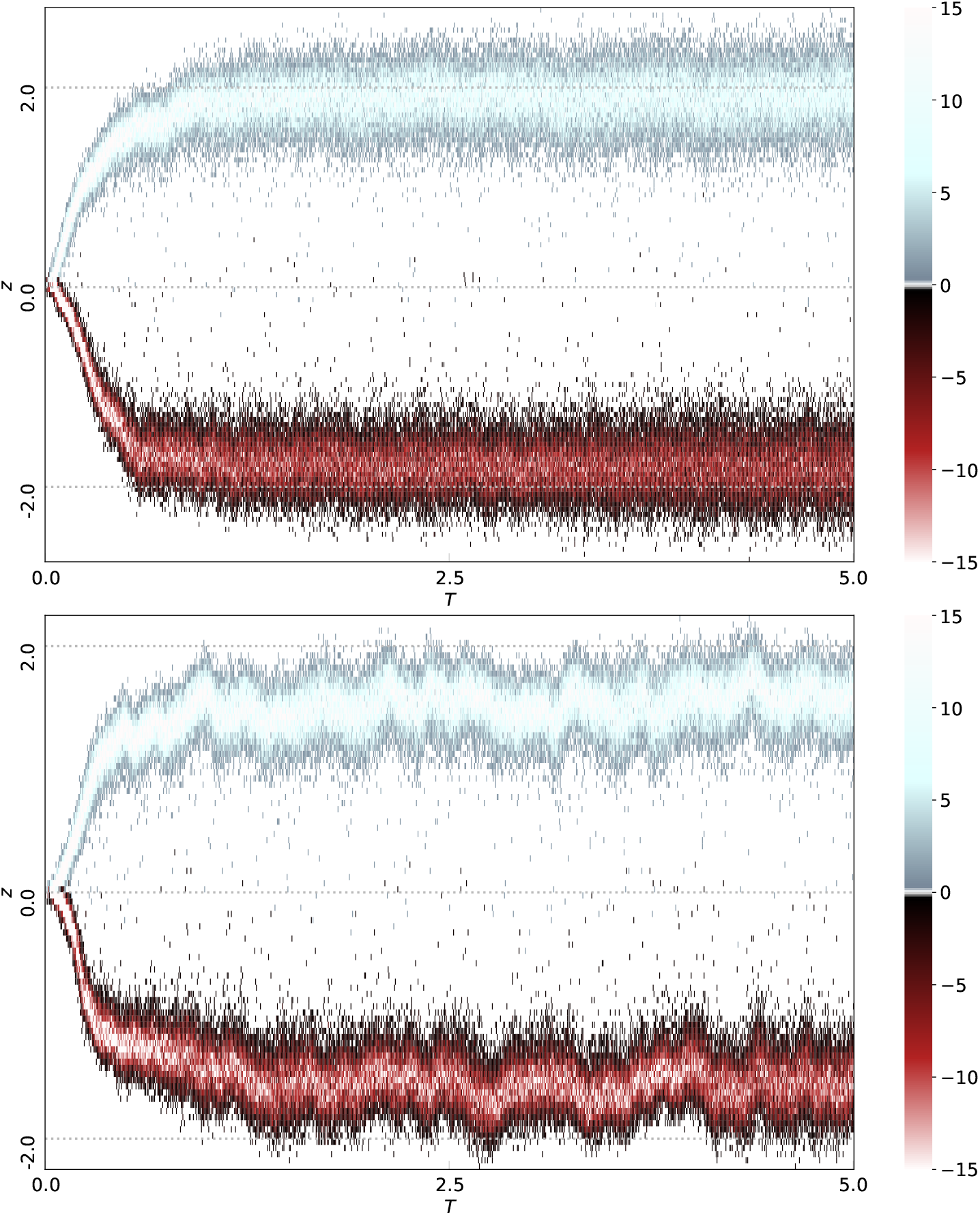
The evolution of individual trait values *z* for a single run for 5 × 2*N*_*e*_ generations. The local optima are represented by the dotted line (*θ*_−_ = −2 and *θ*_+_ = 2). Without (top panel) and with GBMs (*U*_*g*_/*U*_*l*_ = 2, bottom panel) for the “weak-frequent” scenario with *U* = 5 × 10^−3^. Individuals (sample of 100 per population) are binned by phenotype. The shade of each dot represents the number of individuals per bin (see colorbar). Note that the higher mutation rate combined with weak selection induces more variance in the population than in the ‘strong-rare’ scenario.

**Figure S5:**
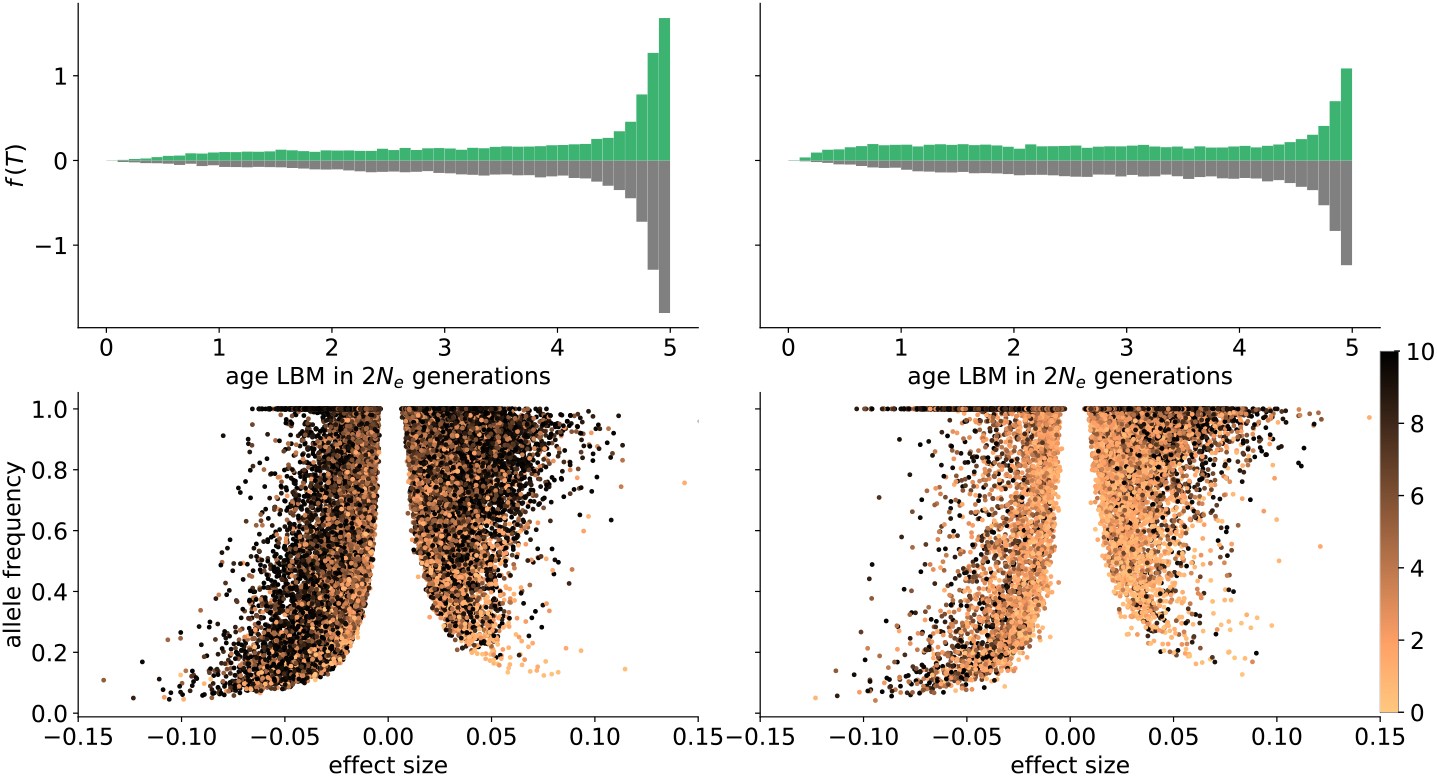
The impact of GBMs on the age distribution of LBMs. Results without GBMs (left) and with GBMs (*U*_*g*_/*U*_*l*_ = 2, right) for “the weak-frequent” scenario, for the population adapting to *θ*_+_, at the dynamic equilibrium phase (5×2*N_e_* generations). Top row: the age distribution of LBMs (across the genome and across 200 replicate runs) weighted by their frequency and effect size. Age is measured in 2*N_e_* generation with 0 representing the time of sampling. The histograms on the top (green) and bottom (grey) correspond to LBMs with positive and negative local effects respectively. Bottom row: LBM effect size plotted against the allele frequency. Young and old LBMs grade from orange to black respectively (see color bar).

**Figure S6:**
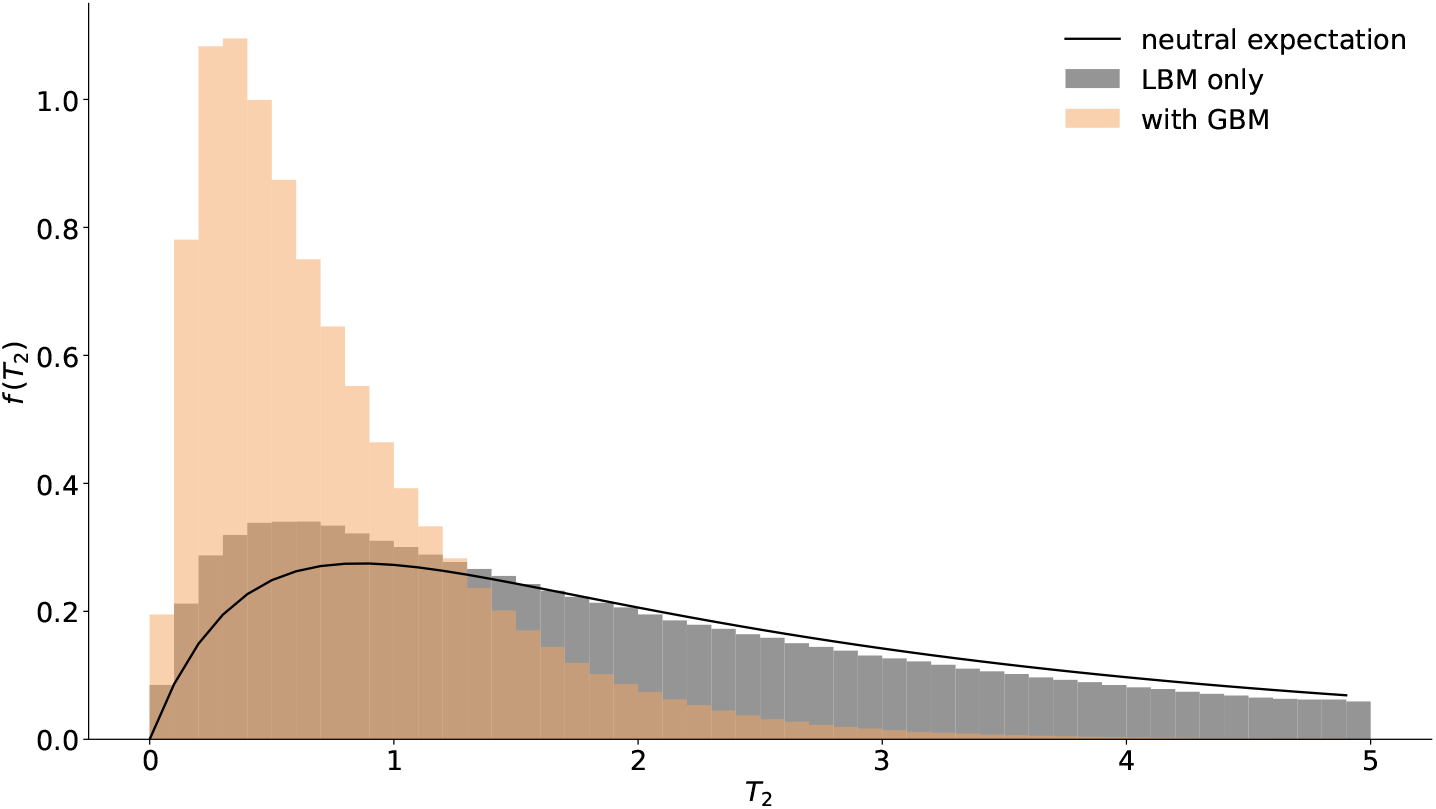
The genome-wide distribution of between population coalescence times *f*(*T*_2_) for the “weak-frequent” scenario. The neutral expectation (assuming *M* = 1) (Lohse et al., 2016, eq. 10) is shown as a gray solid line, distributions. The absence and presence of GBMs (*U*_*g*_/*U*_*l*_ = 2) are shown in gray and orange respectively. 87% and 99% of coalescence times respectively are smaller then 5*N*_*e*_. This data corresponds to figure S3 panel (2,3).

**Figure S7:**
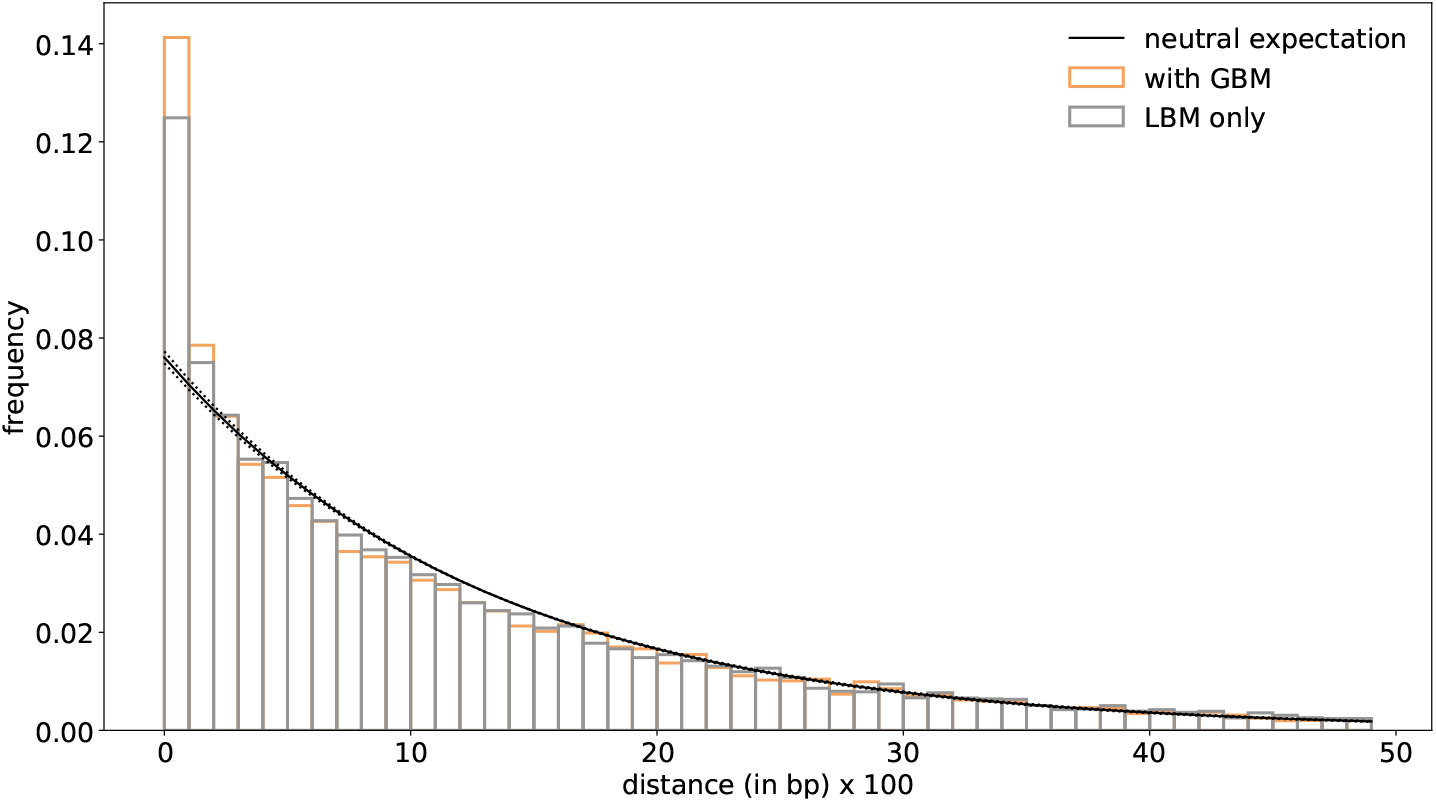
The distribution of pairwise distances between consecutive LBMs contributing positively to divergence (the 75 LBMs of largest effect) for the “weak-frequent” case after 5*N*_*e*_ generations. Frequencies are weighted by the mean contribution of each pair of LBM to local adaption. The black line shows the expected exponential distribution of pairwise distances in the absence of clustering, i.e. assuming loci are distributed uniformly at random (dotted line is 95% confidence interval).

**Figure S8:**
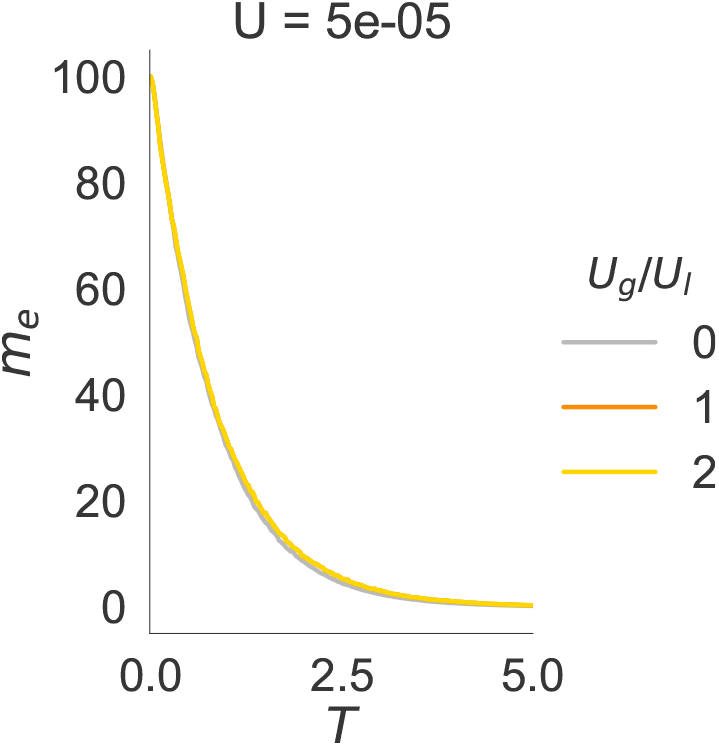
The relative fitness of migrants for the “strong-rare” scenario in the sweep-based model (BU2S) (Flaxman et al., 2013). Time is measured in 2*N*_*e*_ generations. *U*_*g*_/*U*_*l*_ ∈ {0, 1, 2}, gray, yellow and orange respectively.

**Figure S9:**
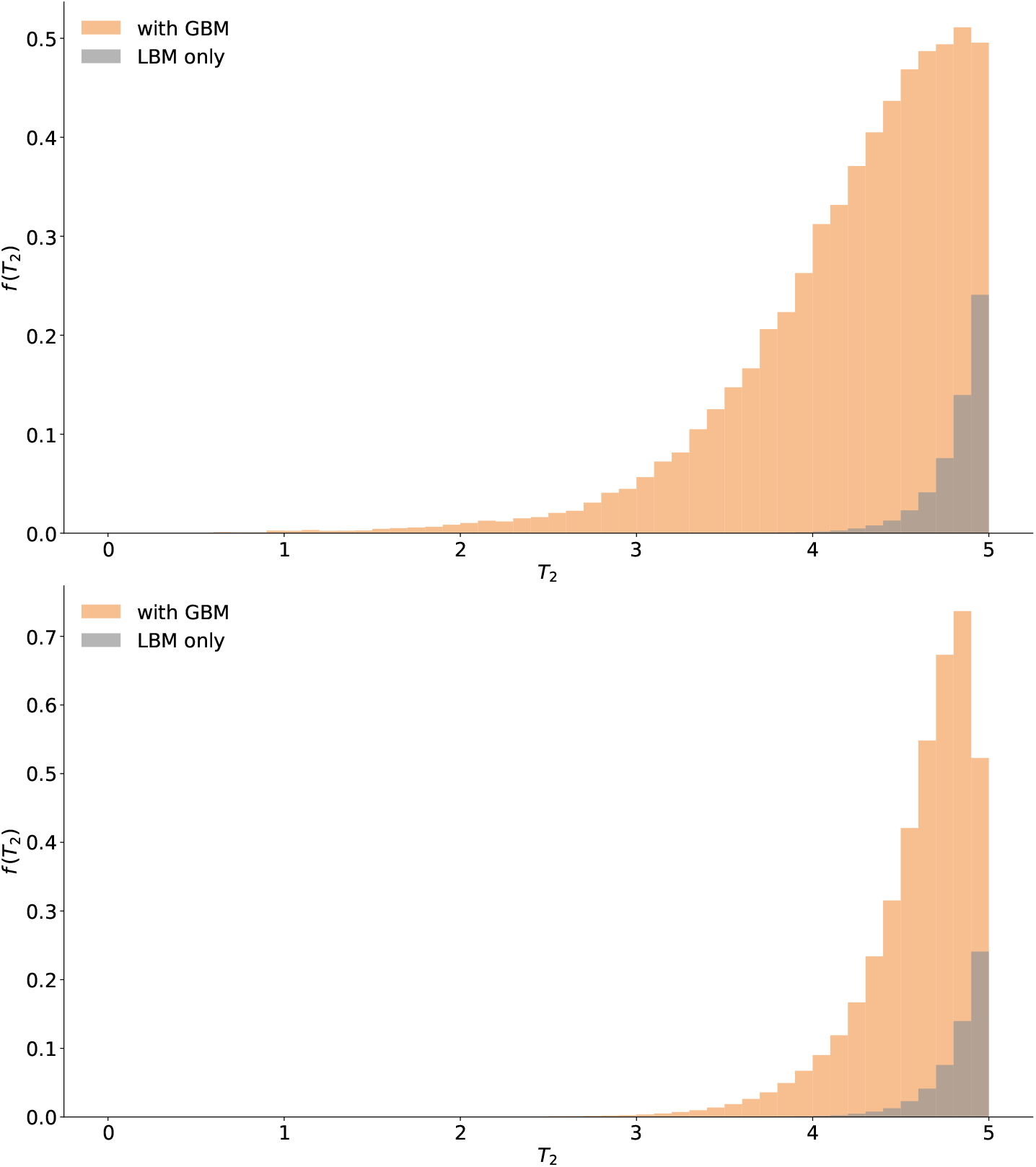
The genome-wide distribution of pairwise coalescence times *f*(*T*_2_) for the sweep-based model (top: “strong-rare”, bottom: “weak-frequent”). Time in 2*Ne*_*e*_ generations, *U*_*g*_/*U*_*l*_ = 2. 0.05% (LBMs only) and 60% (with GBMs) of coalescence times are smaller then 5 (“strong-rare”) and 0.05% (LBMs only) and 40% (with GBMs) for the “weak-frequent” case.

